# Adipose-derived stromal cells preserve pancreatic islet function in a transplantable 3D bioprinted scaffold

**DOI:** 10.1101/2022.05.30.494035

**Authors:** Shadab Abadpour, Essi M. Niemi, Linnea Strid Orrhult, Carolin Hermanns, Rick de Vries, Liebert Parreiras Nogueira, Håvard Jostein Haugen, Dag Josefsen, Stefan Krauss, Aart van Apeldoorn, Paul Gatenholm, Hanne Scholz

## Abstract

Intra-portal islet transplantation is the method of choice for treatment of insulin dependent type 1 diabetes, but its outcome is hindered by limited islet survival due to the immunological and metabolic stress post transplantation. Adipose-derived stromal cells (ASCs) promise to improve significantly the islet micro-environment but an efficient long-term delivery method has not been achieved. We therefore explore the potential of generating ASC enriched islet transplant structure by 3D bioprinting. Here, we fabricate a double-layered 3D bioprinted scaffold for islets and ASCs by using alginate-nanofibrillated cellulose bioink. We demonstrate the diffusion properties of the scaffold and report that human ASCs increase the islet viability, preserve the endocrine function, and reduce pro-inflammatory cytokines secretion *in vitro*. Intraperitoneal implantation of the ASCs and islets in 3D bioprinted scaffold improve the long-term function of islets in diabetic mice. Our data reveals an important role for ASCs on the islet micro-environment. We suggest a novel cell therapy approach of ASCs combined with islets in a 3D structure with a potential for clinical beta cell replacement therapies at extrahepatic sites.

## Introduction

Cell-based therapies represent a game-changer to exogenous insulin treatment of type 1 diabetes (T1D). Today, intraportal islet transplantation is the standard care in many countries with the ability to improve glycaemic control and decrease hypoglycaemic episodes for patients with severe T1D [1, 2]. Despite this, the islet transplantation procedure is limited by several confounding factors including inadequate vascularisation and deleterious islet responses to an inflammatory and immunogenic micro-environment. This leads to loss of islets, poor engraftment and decline in islet function [3–5]. Delivery of islets in hydrogel-based encapsulation scaffolds to animals and human have been explored over the years and reported to protect islets from immune rejection [6]. However, the strategy comes with a price of decreased islet survival, loss of islet function and limited clinical outcomes. This is mainly due to the reduced supply of oxygen and nutrients to the encapsulated islets [7–10]. One solution for more optimal islet survival is open scaffolds applying 3 dimensional (3D) bioprinting technology which allows better tissue integration and enables transplantation to more accessible extrahepatic sites such as subcutaneous and intraperitoneal [6, 11, 12].

Indeed, 3D bioprinting technology is being used for several cell delivery applications [13–17]. The technology allows precise control of scaffold architecture. Moreover, the control over homologous cell distribution within the scaffolds and the possibility to use multiple cell types at once allow fabrication of complex 3D cell delivery systems [18]. Alginate has been tested in various 3D bioprinting applications because of its high biocompatibility and low cytotoxicity [19]. Despite the positive impact of alginate on cells, its low viscosity hinders printing fidelity, which prevents shape retention of bioprinted scaffolds after deposition from the printer nozzle [20]. Cellulose is a natural biopolymer and has been suggested as a potential biomedical material with high degree of shear thinning properties and biocompatibility [21]. Nanofibrillated cellulose (NFC) combined with alginate as a bioink has been investigated previously to generate more stable 3D bioprinted structures including cartilage structures such as components of the human ear pinna and sheep meniscus [22–26]. Recently, alginate-based 3D bioprinting has been applied for pancreatic islets and stem cell-derived insulin-producing cells [27, 28]. Marchioli et al. demonstrated the proof of principle for bioplotting primary islets with alginate/gelatin mixture as bioink [29]. Duin et al. successfully generated pancreatic islet devices by 3D bioprinting alginate/methylecellulose scaffolds [28]. However, these studies reported poor functionality in regards to glucose sensing and insulin secretion [28, 29]. Islets during and post isolation get exposed to changes in their micro-environment [30]. These changes can induce stress-related cell signalling. Additionally, bioprinting islets to scaffolds create a barrier for the islets which might delay or hinder glucose sensing and insulin secretion. These factors potentially reduce the function and viability of islet scaffolds post printing. To overcome these challenges, one strategy is to support islets post isolation and bioprinting procedures by combining islets with supporting cells to prolong the islet function.

Human ASCs can be harvested safely from adipose tissue by lipoaspiration [31]. The molecular properties of ASCs exhibit several favorable features for supporting islet cells. In particular, the ASC secretome has been reported to contain high levels of vascular and intracellular adhesion molecules as well as a diverse range of cytokines and growth factors [32, 33]. The paracrine effects of ASCs have been investigated on various cell-based therapies and wound healing over the years [34, 35]. Moreover, co-culture and co-transplantation of syngeneic ASCs with marginal allogeneic islet mass showed prolonged portal graft survival, islet function and glucose tolerance in diabetic immunocompetent mice [33, 36, 37]. In these experiments, the paracrine effects of ASCs through secretion of protective and anti-apoptotic factors have been suggested to be central on improved islet function and viability [38, 39].

In this study, we fabricated a 3D bioprinted double-layered alginate/NFC scaffold combining mouse or human islets with human ASCs. We show an even distribution of islets and ASCs in each of the scaffolds, and demonstrate the diffusion of small molecules with various sizes through the biomaterial. Our data demonstrate the supportive role of ASCs on the function and viability of mouse or human islets in the alginate/NFC scaffolds *in vitro*. This is accompanied by a reduction in the secretion of selected pro-inflammatory cytokines by the islets. Mouse Islets together with human ASCs in 3D bioprinted scaffolds preserve islet function implanted at the intraperitoneal (IP) site of diabetic mouse model up to 60 day post implantation. Implantation of scaffolds with minimal mass of human islets at the IP site of the same mouse model showed scaffold function up to 30 days post implantation.

## Results

### Fabrication of the double-layered islet scaffold and the study outline

To develop a multicellular scaffold combining pancreatic islets and ASCs, we designed a scaffold with two layers. The bottom layer (shown as blue) for ASCs and an orthogonal grid as the top layer (shown as pink) for mouse or human islets (Fig 1). Using a bioink composed of NFC and alginate (alginate/NFC scaffold; Cellink AB), we first investigated the diffusion properties of the biomaterial. We then bioprinted alginate/NFC scaffolds containing mouse islets and human ASCs (mIslet+hASC), mouse islets alone (mIslet-alone), human islets and human ASCs (hIslet+hASC) and human islets alone (hIslet-alone) and studied the distribution of islets within scaffolds, viability and islet function *in vitro* (Fig 1). Subsequently, mouse or human islet scaffolds were implanted to the IP site of immunocompromised diabetic mice (Fig. 1).

**Fig 1:**
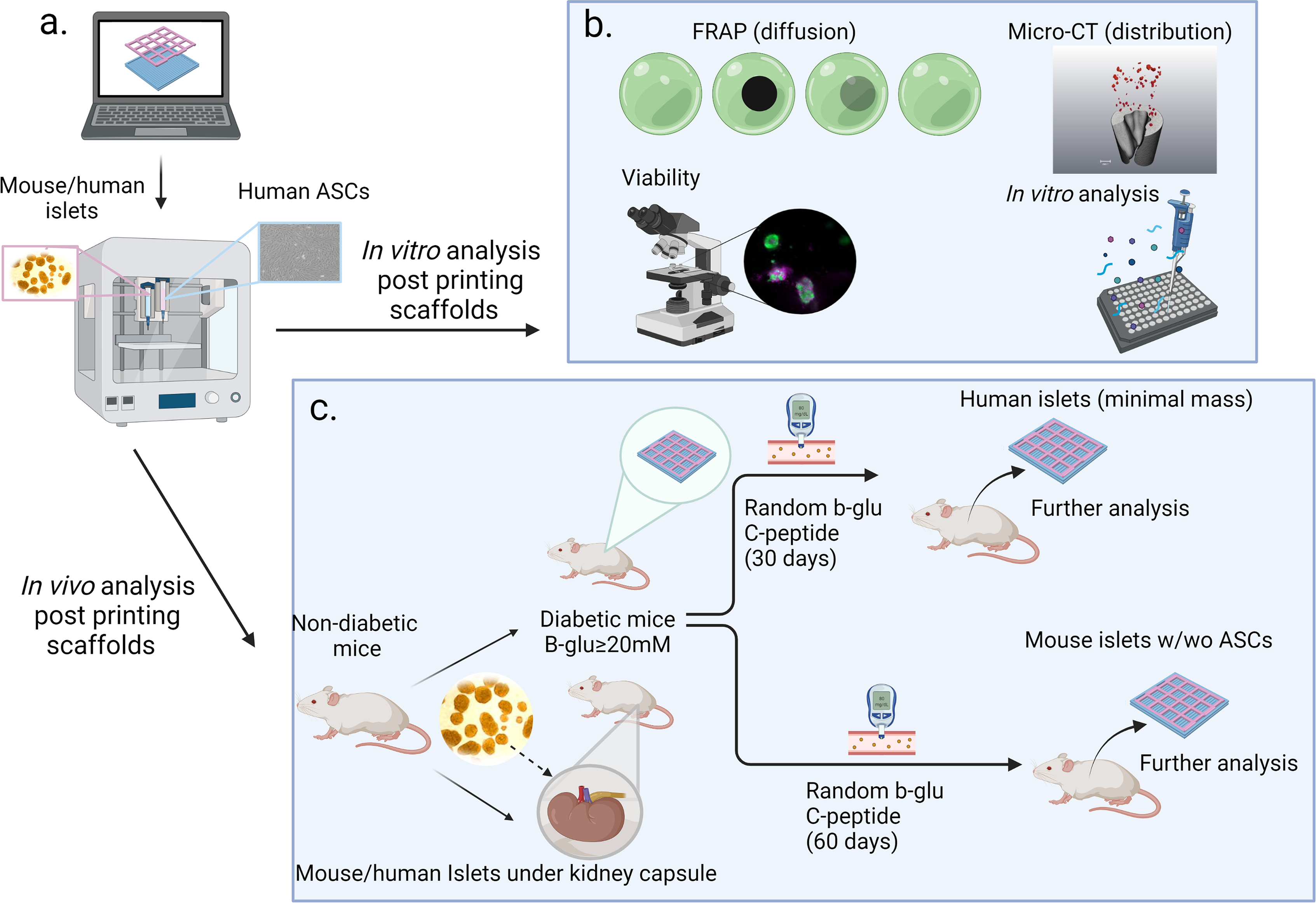
Study outline: 3D bioprinted alginate/NFC scaffolds containing islets and ASCs were fabricated and the scaffold function was investigated by using both *in vitro* and *in vivo* analysis. After the 3D bioprinting procedure (a), the diffusion properties of our bioink were investigated by Fluorescent Recovery After Photobleaching (FRAP) technology (b). The number of islets per scaffold and the distribution of islets were analysed using micro-computated tomography (micro-CT) imaging (b). The function and viability of scaffolds were studied post printing (b). For the *in vivo* analysis, the scaffolds containing human or mouse islets with/without (w/wo) ASCs were implanted to the IP site of the immunocompromised mouse model and followed up for 30 days or 60 days respectively, before scaffolds were explanted for further immunofluorescent analysis (c).

### Alginate/NFC scaffolds show diffusion to various sizes of dextran molecules

To mimic the diffusion properties of alginate/NFC bioink, FRAP imaging was used to assess the diffusion behaviour of fluorescent labelled dextran molecules with sizes ranging from 3-70 kDa. A small area of the gel (Ø60 μm) was imaged (Fig. 2a, Video supplement 1-3), after the area was bleached (Fig. 2b). Dextran molecules were allowed to diffuse freely back to the bleached area, leading to a recovery of the fluorescent signal over time (Fig 2 c-d). This was quantified and normalized to the fluorescence intensity before bleaching (Fig. 2e). The evaluation of the apparent diffusion constant for the alginate/NFC scaffolds incubated with different sizes of dextran molecules showed a significant increase in diffusion properties of scaffolds incubated with 3-5 kDa compared to scaffolds incubated with 20 kDa and 70 kDa (Fig. 2f). Therefore, our data reported a difference in diffusion properties of molecules with different sizes. However, all analysed dextran molecules could diffuse though the bioprinted scaffolds confirming the exchange of molecules between 3-70 kDa in size.

**Fig 2.**
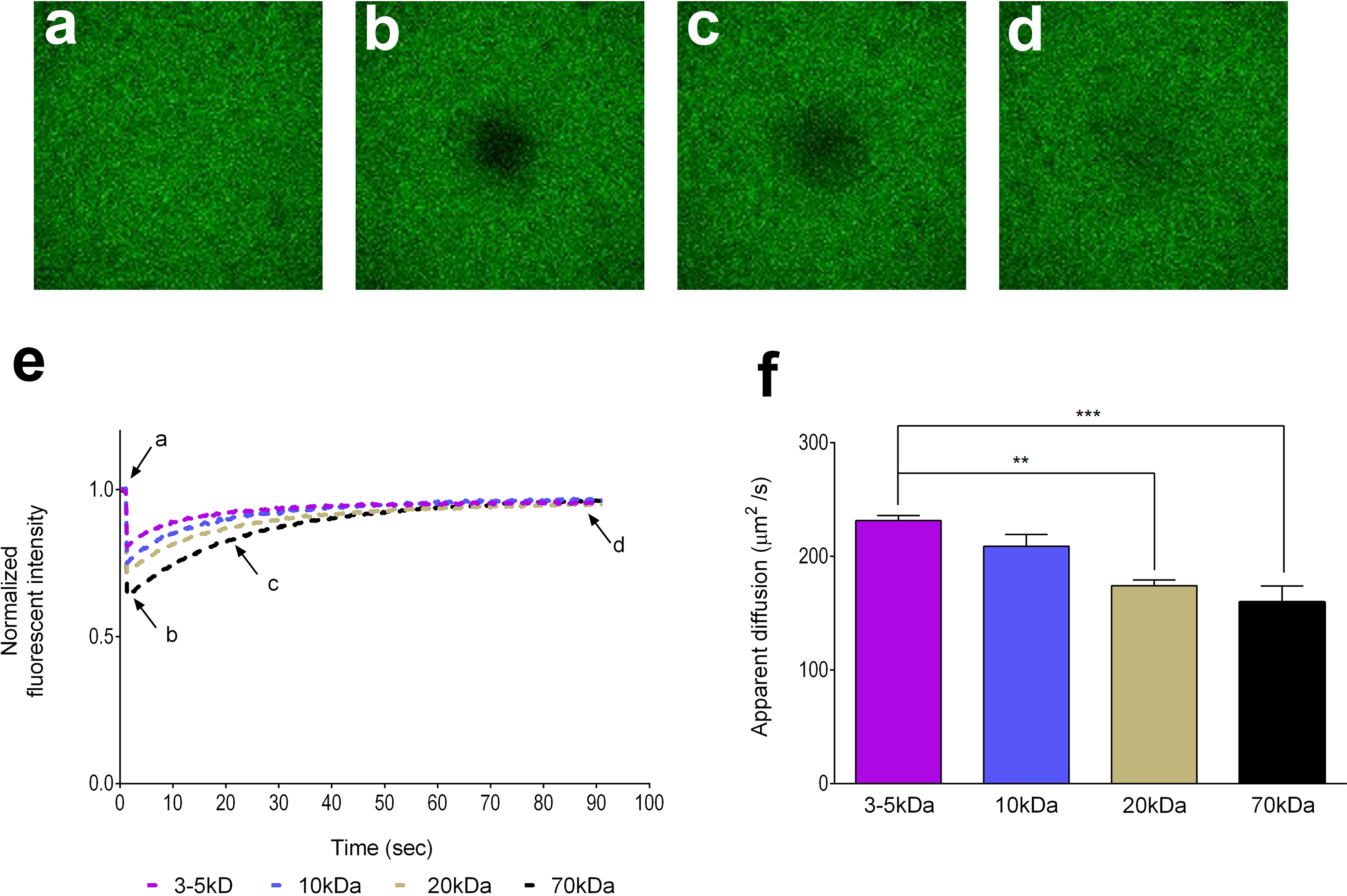
with 3 supplements (ESV 1-3). Diffusion of dextran molecules with 3-70 kDa in size to the bioprinted alginate/NFC scaffolds. Confocal images of the alginate/NFC scaffolds that were saturated with fluorescent-labelled dextran molecules before bleaching (a), during bleaching (b), in early-recovery phase (c) and in late-recovery phase (d). The fluorescence recovery curve obtained by determining fluorescence intensity of the bleached area (e), highlighting the moments when images a-d were taken. Obtained fluorescence recovery curves were subsequently used to calculate the apparent diffusion properties of fluorescent-labelled dextran molecules through the scaffolds (f). All scaffolds were measured five times at different locations (technical replication). In all analyses, data are presented as mean±SD and analysed by one-way ANOVA with Bonferroni corrections and Mann–Whitney U test. **p < 0.01 and ***p < 0.001 vs. 3-5kDa group.

### ASCs improve the function and viability of mouse islets in 3D bioprinted alginate/NFC scaffold

To investigate the effects of human ASCs on mouse islet function in the scaffold, we performed glucose stimulated insulin secretion (GSIS) on days 1, 8 and 14 post printing. Insulin secretion in response to 20mM glucose (stimulated levels) was increased in the mIslet-alone group; however, islets in this group failed to reduce insulin secretion to basal levels after transferring the scaffolds from the solution containing 20mM glucose to the solution containing 1.67mM (basal levels) (Fig. 3a). mIslet+hASC scaffolds revealed improvement in islet response to different levels of glucose (Fig. 3a). This response was similar to the free-floating mouse islets from the same batch of printed islets (Fig. 3a). Consequently, the stimulation index (SI) was also improved in mIslet+hASC group compared to the mIslet-alone group (Day 1, mIslet+hASC 2.581 ± 0.14 vs. mIslet-alone 1.108 ± 0.04, p<0.05, Day 8, mIslet+hASC, 1.108 ± 0.04 vs. mIslet-alone 1.575 ± 0.05, Day 14, mIslet+hASC 1.410 ± 0.06329 vs. mIslet-alone 0.7989 ± 0.1095) (Fig. 3b).

**Fig 3:**
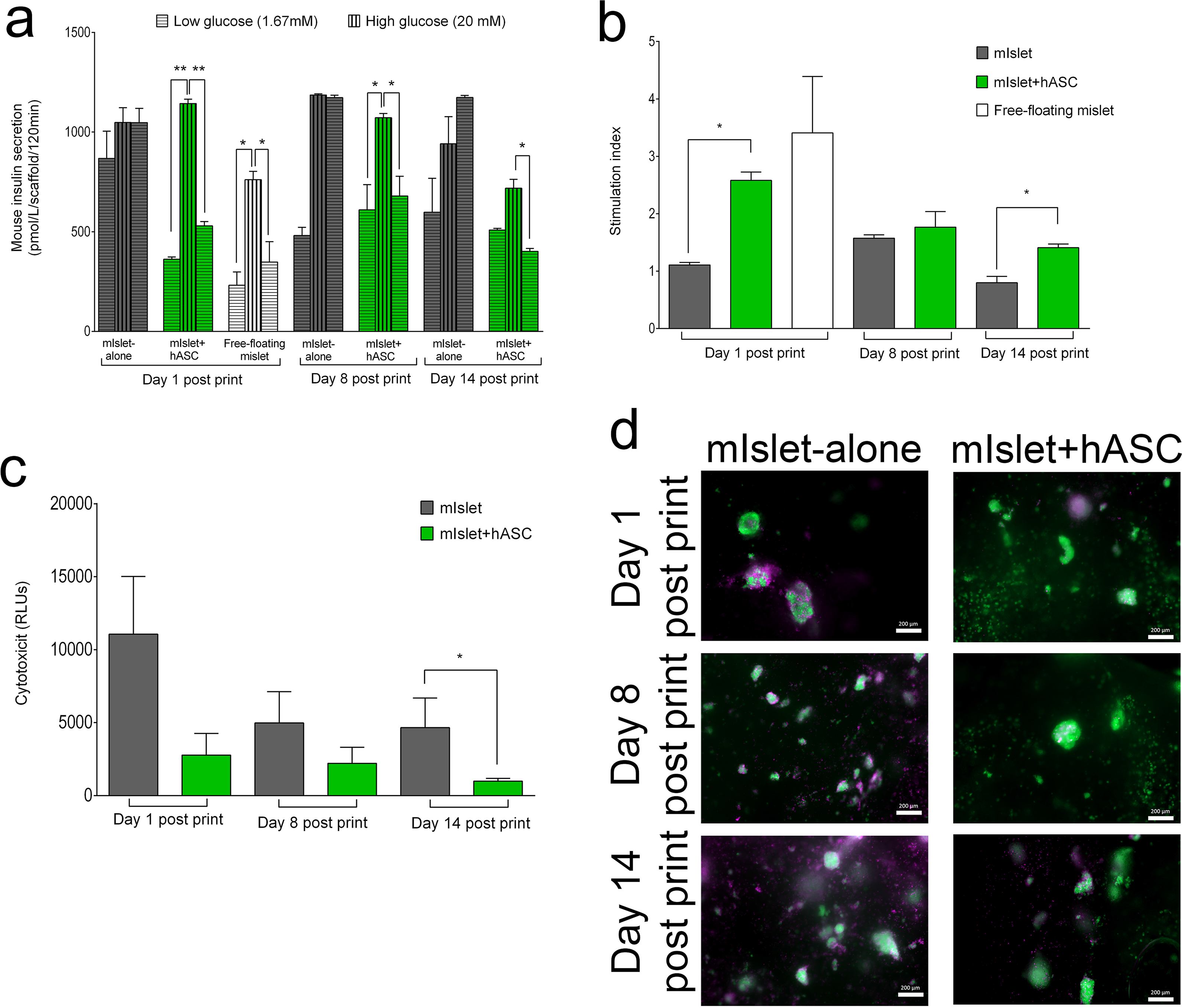
mIslet+hASC scaffolds showed improvement in insulin secretion in response to glucose and cell viability. (a) Insulin secretion in response to basal (1.67 mmol/l) and stimulated (20 mmol/l) levels of glucose measured by ELISA (b) and calculated as stimulation index for mouse islets w/wo ASCs analysed on days 1, 8 and 14 post print (a-b). Cytotoxicity was measured by analysing the levels of secreted ADK enzyme released by mouse islets w/wo ASCs in alginate/NFC 3D bioprinted scaffolds, analysed on days 1, 8 and 14 post print (c). FDA (live cell-green) / PI (dead cell-purple) staining followed up by visualization of mouse islets w/wo ASCs, taken on days 1, 8 and 14 post print (d). Magnification 10x, scale bar 200 µm. n = 4 independent biological replicates. In all analysis, data are presented as mean±SD and analysed by one-way ANOVA with Bonferroni corrections and Mann–Whitney U test. *p < 0.05 and **p < 0.01 vs. high glucose (20mM) group, * p<0.05 vs. mIslet-alone group.

Release of the mitochondrial Adenylate Kinase (ADK) enzyme is well-known as a sign of cell death [40]. the mIslet+hASC group showed a trend in reducing levels of secreted ADK compared to the mIslet-alone group on days 1, 8 post print (Fig. 3c). We found a significant difference in the levels of ADK on day 14 post print in this group compared to the mIslet-alone group (Fig. 3c). In addition, mIslet+hASC group showed less (propidium iodide) PI-positive cells analysed by live/dead immunofluorescent staining of cells with (fluorescein diacetate) FDA/PI, which supported our ADK data and shows improved viability in mIslet+hASC group compared to the mIslet-alone group (Fig. 3d).

### ASCs reduce levels of secreted pro-inflammatory cytokines in 3D bioprinted mouse islet scaffolds *in vitro*

Three pro-inflammatory cytokines that have shown to be secreted by islets upon transplantation were selected for analysing the impact of ASCs on islets [41–43]. In earlier work, co-culture of human ASCs and human islets has shown to reduce secretion of these cytokines previously [32]. In our setting, mIslet-alone scaffolds showed an increase in the levels of secreted mouse specific pro-inflammatory cytokines, (MCP-1/ known as CCL2), interferon gamma-induced protein-10 (IP-10/known as CXCL10) and growth-regulated oncogene-α (GRO-α/known as CXCL1) (Fig 4a-c) measured on days 1, 8 and 14 post print. In contrast, presence of human ASCs in the mIslet+hASC scaffolds reduced significantly the secretion of all three cytokines significantly on day 1 post print. Our data showed also a significant reduction of MCP-1 on day 8 post print, and GRO-α and IP-10 on day 14 post print in mIslet+hASC group compared to the mIslet-alone group (Fig 4a-c) (MCP-1, Day 1: mIslet+hASC 2.87 ± 1.44 pg/ml vs. mIslet-alone 19.09 ± 1.04 pg/ml, p<0.001, Day 8, mIslet+hASC, 8.28 ± 2.87 pg/ml vs. mIslet-alone 20.87 ± 3.59 pg/ml, p<0.05 – IP-10, Day 1, mIslet+hASC 2.532 ± 0.98 pg/ml vs. mIslet-alone 6.538 ± 1.51 pg/ml, p<0.05, Day 8, mIslet+hASC 1.20 ± 1.20 pg/ml vs. mIslet-alone 1.89 ± 1.89 pg/ml – GRO-α, Day 1, mIslet+hASC 70.14 ± 12.75 pg/ml vs. mIslet-alone 559.3 ± 113.5 pg/ml, p<0.01, Day 8, mIslet+hASC 37.57 ± 10.43 pg/ml vs. mIslet-alone 83.21 ± 15.78 pg/ml). The data show that the presence of human ASCs in islet scaffolds protect inflammatory status of mouse islets up to 2 weeks post print.

**Fig 4:**
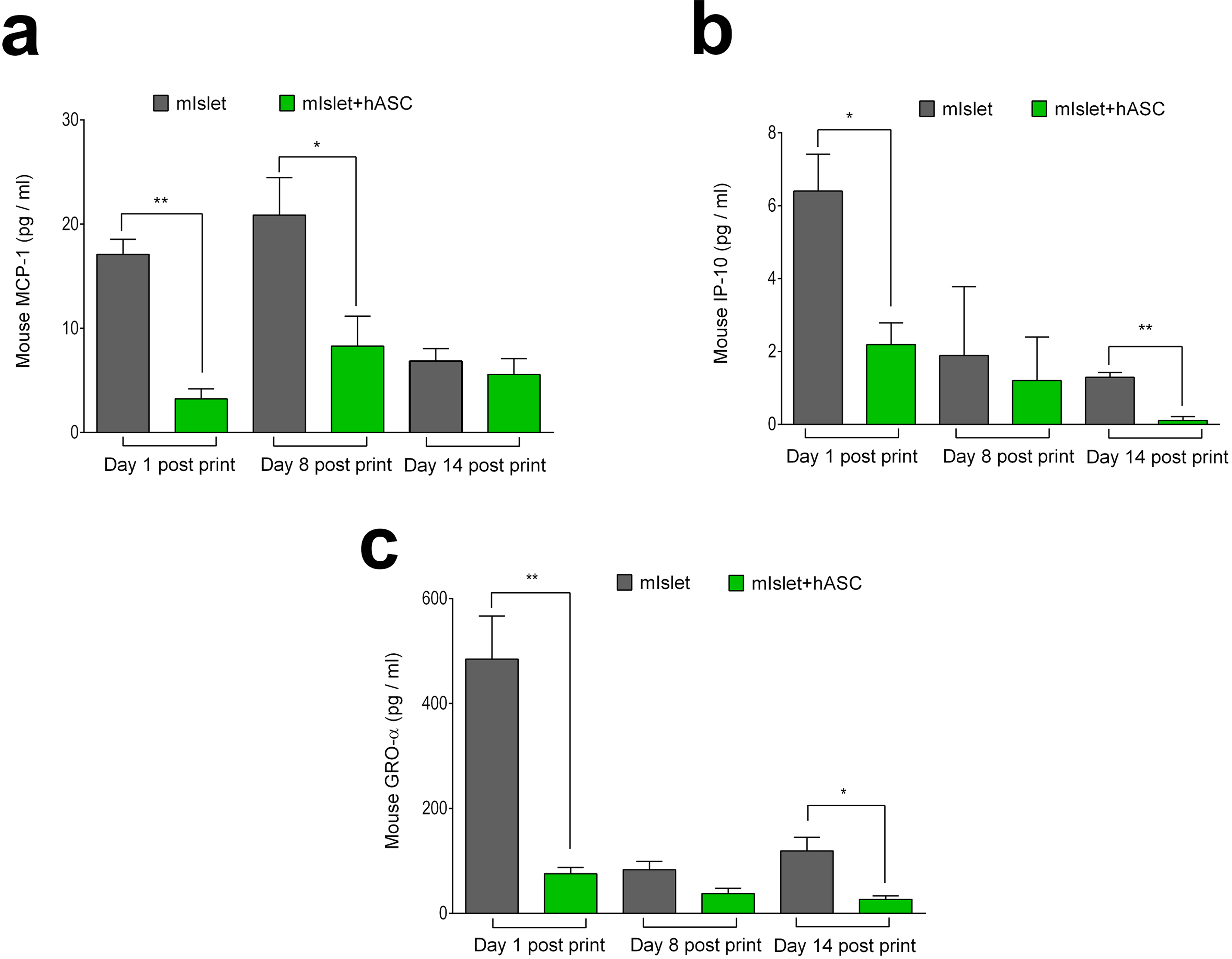
mIslet+hASC scaffolds improved inflammatory status of the printed islets. Assessment of secreted mouse proinflammatory cytokines MCP-1 (a), IP-10 (b) and GRO-α (c) by multiplex assay in cell-free supernatant collected from the bioprinted scaffolds with mouse islets w/wo ASCs on days 1, 8 and 14 post print. In all analysis, data is presented as mean±SD and analysed by Mann Whitney U-test. n= 4 independent biological replicates. * p<0.05, and ** p<0.01vs. mIslet-alone group.

### Human islets were evenly distributed in 3D bioprinted scaffolds and ASCs improved human islet function and viability

Next, approximately 100 islets with islet size between 100µm to 200 µm were printed (Fig 5a-b, Method supplement, Table supplement 1). Micro-CT imaging of the human islet containing alginate/NFC scaffolds revealed an even distribution within the 3D bioprinted scaffolds (Fig. 5a, Method supplement, Figure supplement 1, Video supplement 4). Insulin secretion in response to stimulated glucose levels was increased in hIslet-alone group; however, these scaffolds showed blunted insulin secretion upon transferring from stimulated glucose to basal glucose levels (Fig. 5c). In contrast, hIslet+hASC group showed improvement in islet response to stimulated and basal glucose levels (Fig. 5c). Free-floating human islets showed the same response to basal and stimulated glucose levels as hIslet+hASC group. Consequently, hIslet+hASC group revealed higher levels of SI compared to the hIslet-alone group (Day 1, hIslet+hASC 1.835 ± 0.22 vs. hIslet-alone 1.239 ± 0.27, Day 8, hIslet+hASC, 2.408 ± 0.42 vs. hIslet-alone 0.7203 ± 0.13, Day 14, hIslet+hASC 2.315 ± 0.38 vs. hIslet-alone 0.7203 ± 0.13) (Fig. 5d). Analysis of the cell viability in 3D bioprinted scaffolds revealed reduction in secreted ADK enzyme in hIslet+hASC group on days 1, 8 and 14 post print compared to the hIslet-alone group (Fig. 5e). In addition, we observed less PI-positive cells in hIslet+hASC group compared to hIslet-alone scaffolds, analysed by FDA/PI staining. This data supports our ADK measurement in the scaffold culture medium and confirm the viability improvement of hIslet+hASC group compared to the hIslet-alone group (Fig. 5e-f).

**Fig 5:**
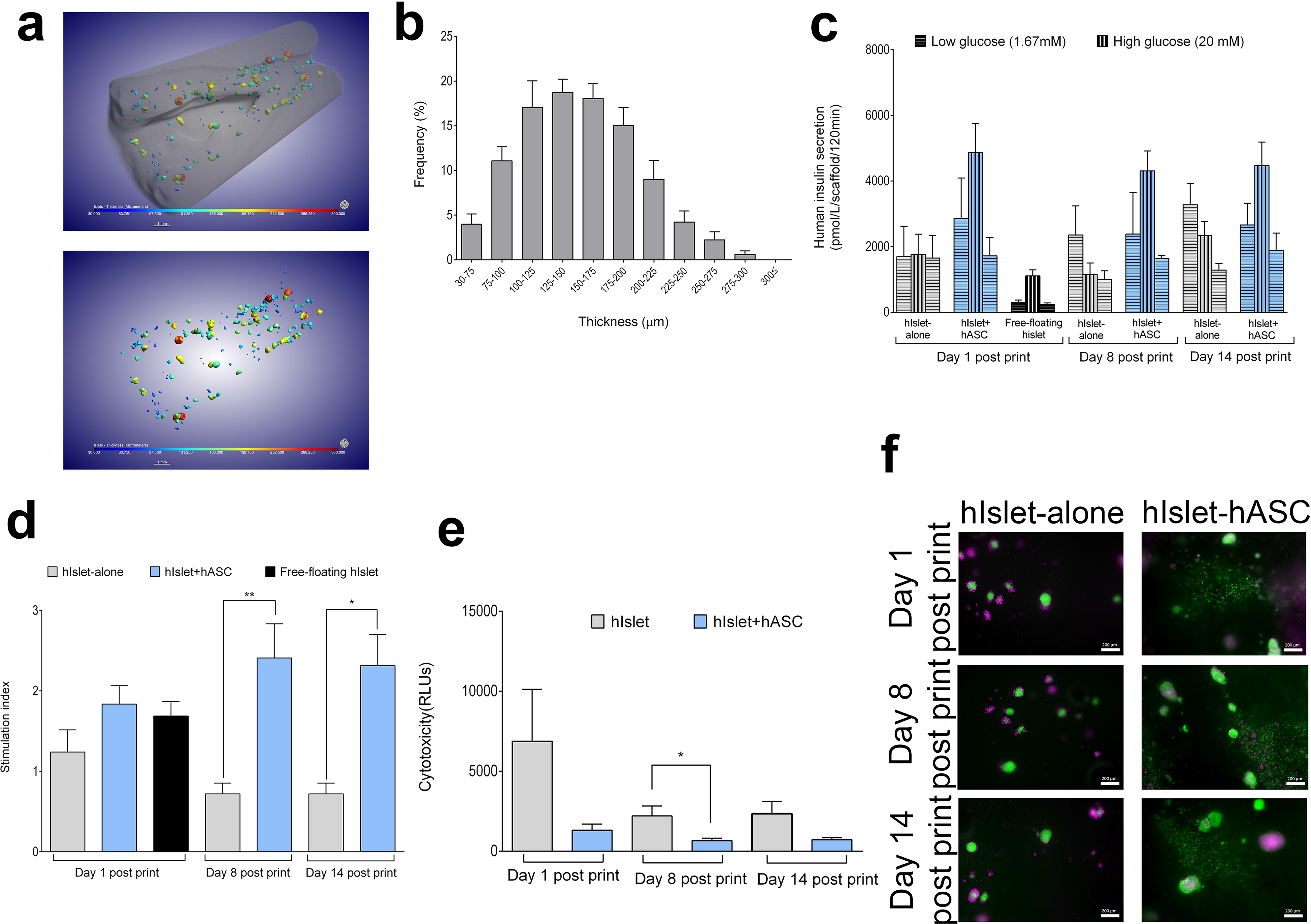
with 3 supplements (ESM-table1, ESR Fig1, ESV 4), Micro-CT imaging of human islets in 3D bioprinted scaffolds and improvement of islet viability and function in hIslet+hASC scaffolds. Micro-CT analysis shows the total number of islets per scaffold (a) and the frequency of islet sizes in 3D bioprinted scaffolds (b). Insulin secretion in response to basal (1.67 mmol/l) and stimulated (20 mmol/l) levels of glucose measured by ELISA (c) and calculated as SI (d) for human islets w/wo ASCs analysed on days 1, 8 and 14 post print. Cytotoxicity was measured by analysing the levels of secreted ADK enzyme released by human islets w/wo ASCs in 3D bioprinted scaffolds on days 1, 8 and 14 post print (e). FDA (live cell-green) / PI (dead cell-purple) staining followed up by visualization of human islets w/wo ASCs taken on days 1, 8 and 14 post print (f). Magnification 10x, scale bar 200 µm. n = 4 independent biological replicates. In all analysis, data are presented as mean±SD and analysed by one-way ANOVA with Bonferroni corrections and Mann–Whitney U test. *p < 0.05 and **p < 0.01 vs. hIslet-alone group.

### ASCs reduce levels of secreted pro-inflammatory cytokines in 3D bioprinted human islet scaffold

The levels of secreted human pro-inflammatory cytokines, MCP-1 (Fig. 6a), GRO-α (Fig. 6b) and IP-10 (Fig. 6c) were measured from the scaffold culture medium on days 1, 8 and 14 post print, which revealed a significant reduction in secretion of MCP-1 and GRO-α on day 1 post print and IP-10 on day 8 post in hIslet+hASC group compared to the hIslet-alone group (Fig 6 a-c). We also observed a reduction in the levels of all three pro-inflammatory cytokines in hIslet+hASC group compared to hIslet-alone group on day 14 post print, although the differences were not significant. This data suggests an anti-inflammatory and protective effects of ASCs on human islets and confirms our previously presented data on the protective effect of ASCs on 3D bioprinted mouse islet scaffolds (Fig 4).

**Fig 6:**
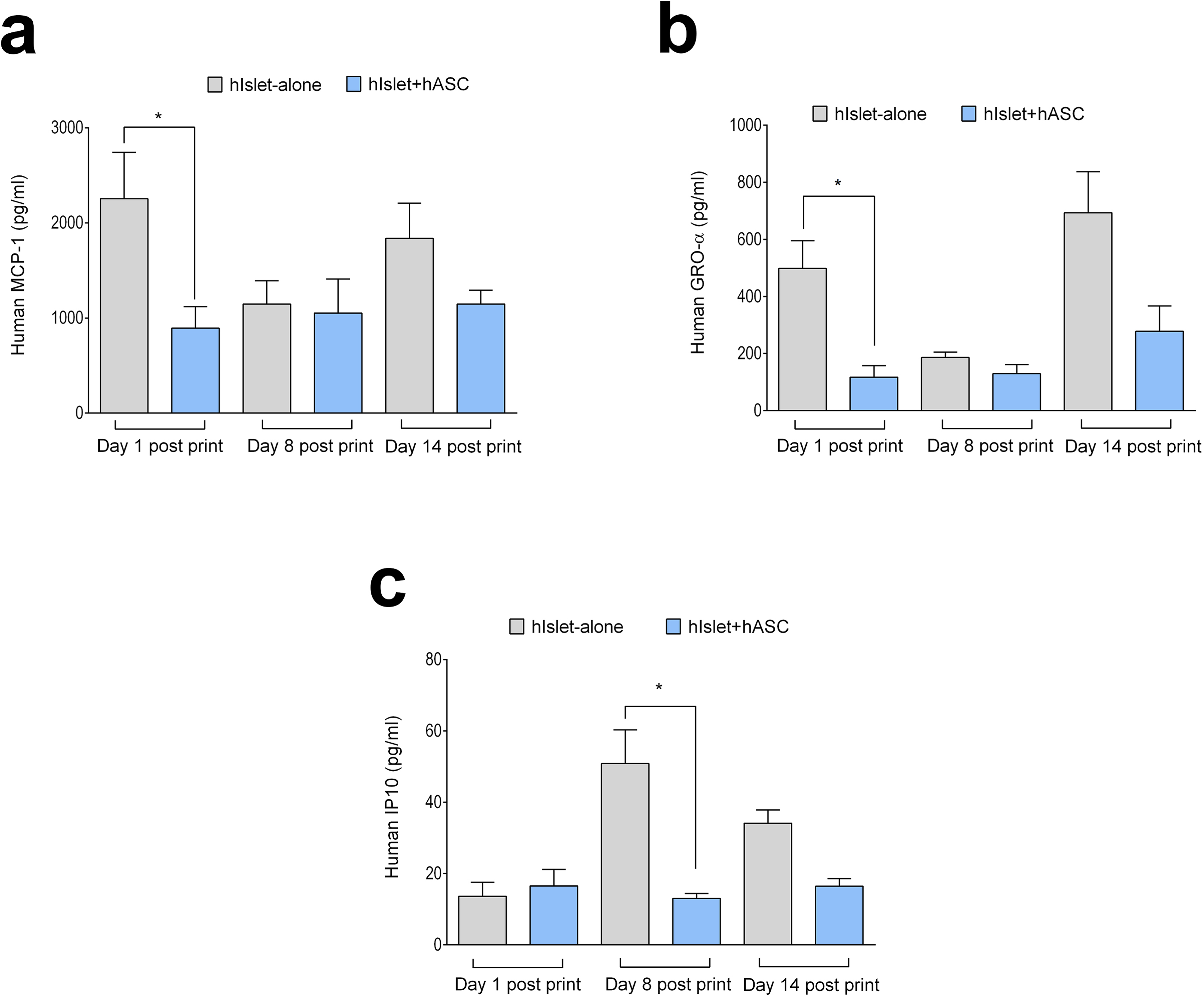
Scaffolds with hIslet+hASC showed reduction in secretion of selected pro-inflammatory cytokines. Assessment of secreted levels of human pro-inflammatory cytokines, MCP-1 (a), GRO-α (b) and IP-10 (c) by multiplex assay in cell-free supernatant collected from hIslet-alone and hIslet+hASC scaffolds on days 1, 8 and 14 post print. In all analysis, data are presented as mean±SD and analysed by Mann Whitney U-test. N= 4 independent biological replicates. * p<0.05 vs. hIslet-alone group.

### Implantation of 3D bioprinted scaffolds containing minimal mass of human islets to the IP site of diabetic mouse model

To investigate the longitudinal adaptation of minimal mass of human islets in 3D bioprinted scaffolds *in vivo,* we fabricated islet scaffolds with 400 human islets (Fig. 7a). 400 human islets have been shown as a non-curative minimal mass graft to investigate human islet function *in vivo* [44, 45]. First, we documented an even distribution of the 400 human islets within the scaffolds by micro-CT imaging (Fig 7a, Method supplement, Table supplement 2, Figure supplement 2 and Video supplement 5). The average size of human islets for this analysis was between 100-200 µm (Fig 7b). Although, implanting scaffolds with minimal mass of human islets (400 human islets) (hIslet-alone) to the IP site of the diabetic mice could not normalize random blood glucose (Fig 7c-d), fasting blood glucose reduced in these animals compared to the diabetic control group (Fig 7e-f). The same results were observed in diabetic mice that received 400 free-floating human islets under the kidney capsule (minimal mass under the kidney capsule) (Fig 7 c-f). 800 human islet grafts is known as the curative full mass of islet graft for treating diabetic mice [45, 46]. Comparing our data on scaffolds or grafts with minimal islet mass with animals received 800 human islets under kidney capsule (full mass under kidney capsule), we observed that the animals with the full islet mass showed a reduction in the levels of random blood glucose and reached euglycemia around day 11 post implantation (Fig 7c-d). We found no significant difference in the levels of fasting blood glucose in animals that received full islet mass compared to the animals that received minimal mass of human islets, either as islet scaffolds or under the kidney capsule (Fig. 7e-f).

**Fig 7.**
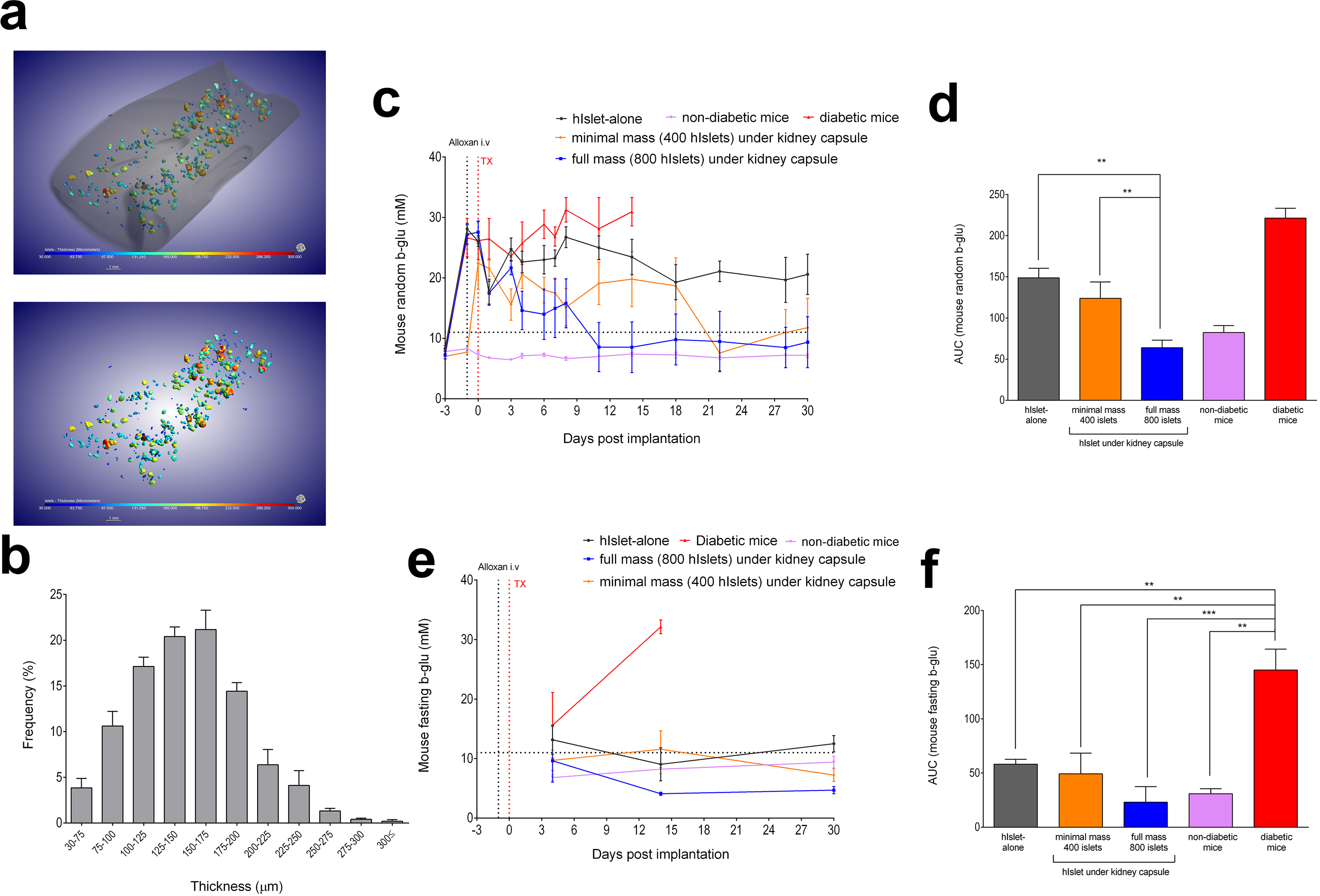
with 3 supplements (ESR Fig 2, ESV 5, ESM table 2): *in vivo* function of scaffolds with minimal mass of human islets implanted to the IP site of the diabetic mice for 30 days post implantation. Micro-CT analysis of the total number of human islets per scaffold (a) and the frequency of islet size in 3D bioprinted scaffolds (b). Random blood glucose measured for 30 days until termination of the study (c) and area under curve (AUC) (d) were calculated for the levels of random blood glucose in mice implanted with scaffolds, islet grafts or control animals. Fasting blood glucose (e) and calculated AUC (f) for mice implanted with islet scaffolds, islet grafts or control animals. In all analysis, data is presented as mean±SD and analysed by one-way ANOVA with Bonferroni corrections and unpaired Mann Whitney U-test. N=3 independent biological replicates and for each replicate 6 mice per group were used. ** p<0.01 vs. mice implanted with full mass of human islets under kidney capsule. ** p<0.01 and *** p<0.001 vs. diabetic mice.

### Human islets in 3D bioprinted alginate/NFC scaffolds remain functional for up to 30 days post implantation to the IP site of diabetic mice

Analysing the functionality of scaffolds with minimal mass of human islets post implantation to the IP site of the diabetic mice revealed secretion of human specific C-peptide from the islets in scaffolds on day 30 post implantation (Fig. 8a). We found no significant difference in the levels of secreted human C-peptide in these animals compared to the mice implanted with minimal mass of human islet grafts under kidney capsule (Fig. 8a). The same results were observed for the ratio of human C-peptide to fasting blood glucose (Fig. 8b). The level of human C-peptide was significantly higher in mice implanted with full mass of human islets under kidney capsule compared to the mice with minimal islet mass either in scaffolds or grafts (Fig 8a). Similarly, we found a significant increase in the ratio of human C-peptide to fasting blood glucose in animals received full mass of islets under kidney capsule compared to the animals with minimal islet mass either as islet scaffolds or as islet grafts (Fig. 8b). At termination of the study on day 30 post implantation, immunofluorescent staining of explanted hIslet-alone scaffolds revealed positive immunofluorescent double-staining of human islets for C-peptide and glucagon, confirming the presence of functional human islets in 3D bioprinted scaffolds 30 days post implantation (Fig. 8c).

**Fig 8:**
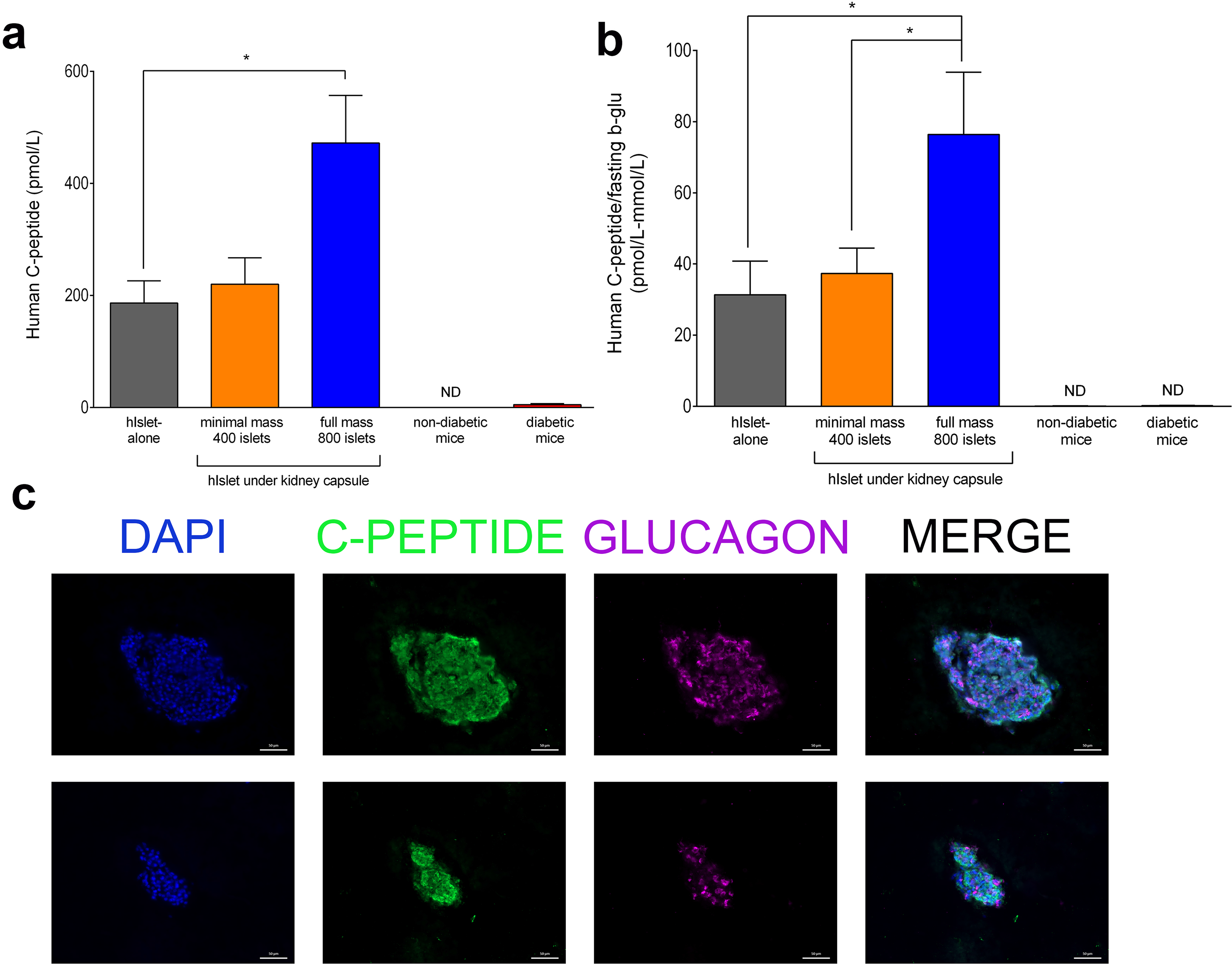
Functional scaffolds with minimal mass of human islets up to 30 days post implantation at the IP site of diabetic mouse model. Circulating human C-peptide (a) and calculated ratio of human C-peptide/fasting blood glucose (b) measured at the termination of studies in mice that received hIslet-alone scaffolds, islet grafts or control animals. Representative immunofluorescent double-staining of explanted human islet scaffolds for C-peptide (green)/glucagon (purple) 30 days post implantation (c). Magnification 10x, scale bar 200 µm. In all analysis, data are presented as mean±SD and analysed by one-way ANOVA with Bonferroni corrections and unpaired Mann Whitney U-test. N=3 independent biological replicates and for each replicate 6 mice per group were used. * p<0.05 and ** p<0.01 vs. mice implanted with full mass of human islets under kidney capsule.

### Mouse islets and human ASCs in 3D bioprinted scaffolds normalized blood glucose in a diabetic mouse model

To assess the effect of human ASCs on mouse islet function and glucose regulation *in vivo,* scaffolds containing 100 mouse islets and human ASCs (mIslet+hASC) were implanted to the IP site of diabetic mice and followed up for 60 days post implantation. All animals implanted with mIslet+hASC showed reduction in random blood glucose to normoglycemic levels up to 7 days post implantation (Fig 9a-c). However, all diabetic animals implanted with only mouse islets (mIslet-alone) scaffolds reached levels of euglycemia 56 days post implantation (Fig 9 a-c). In parallel and to compare our model with well-established *in vivo* model for studying the potency and function of isolated islets, 250 free-floating mouse islets were implanted under the kidney capsule of the diabetic mouse model [47]. Free-floating mouse islets were potent to reduce random blood glucose up to day 2 post implantation (Fig 9a-c). This is similar to animals implanted with mIslet+hASC scaffolds, confirming the positive impact of ASCs on mouse islet function *in vivo.* Measured fasting blood glucose and the analysed AUC post implantation showed a significant reduction in the levels of fasting blood glucose in animals implanted either with mIslet+hASC or mIslet-alone scaffolds compared to the diabetic mice (Fig 9d-e). No significant differences were observed in the levels of fasting blood glucose in diabetic mice implanted with mIslet+hASC scaffolds, mIslet-alone scaffolds or mouse islets under kidney capsule (Fig 9d-e).

**Fig 9:**
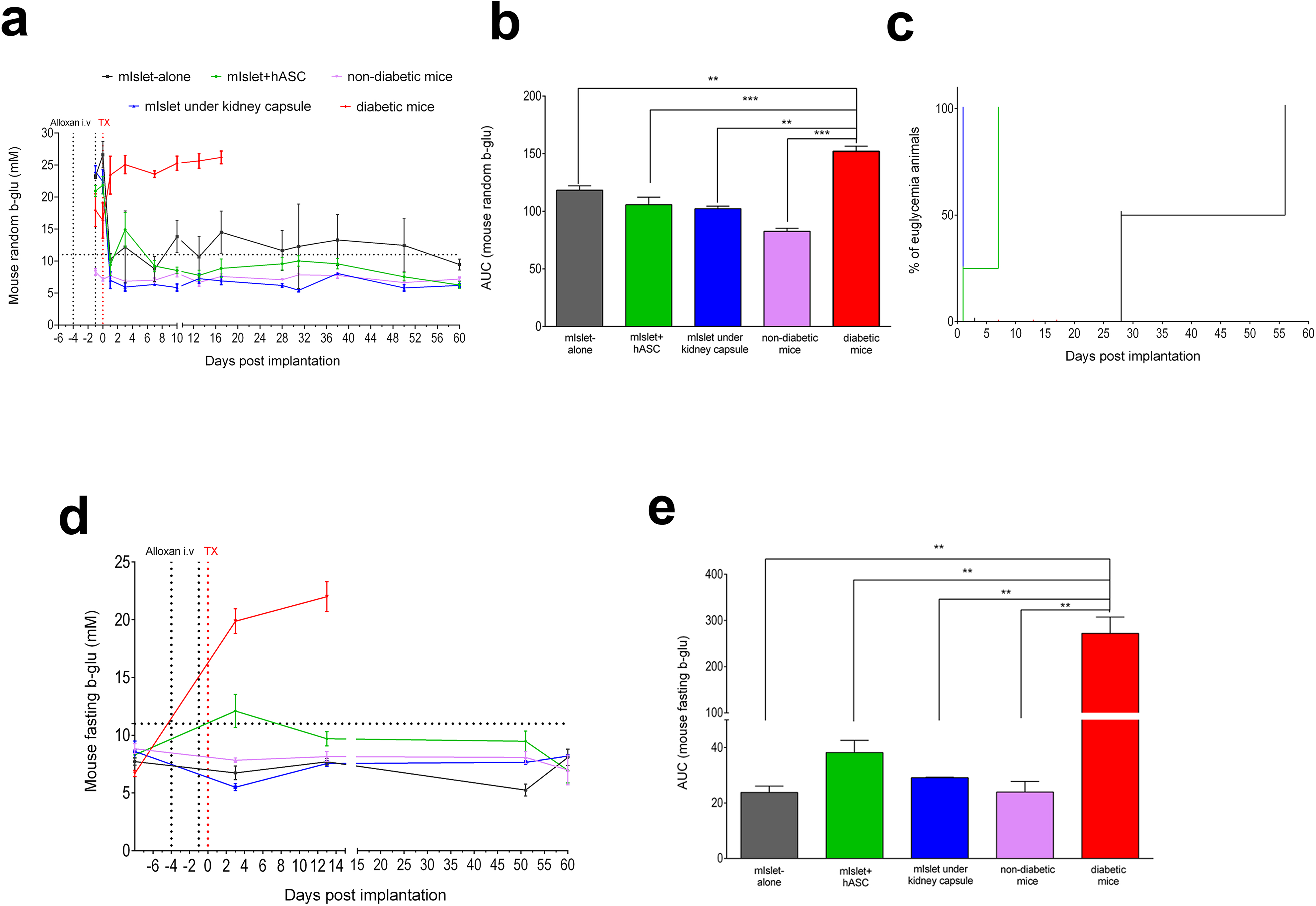
Diabetic mice implanted with mIslet+hASC scaffolds reached normoglycemia 7 days post implantation with reduced levels of fasting blood glucose. Random blood glucose measured until termination of the studies (a) and calculated AUC (b) for the level of random blood glucose in mice implanted with mIslet+hASC or mIslet-alone scaffolds, islet grafts or control animals. Survival curve for mice implanted with scaffolds or islet grafts under kidney capsule (c). Fasting blood glucose (d) and calculated AUC (e) for mice implanted with scaffolds, islet grafts or control animals. In all analysis, data is presented as mean±SD and analysed by one-way ANOVA with Bonferroni corrections and unpaired Mann Whitney U-test. N=3 independent biological replicates and for each replicate 6 mice per group were used. ** p<0.01 and *** p<0.001 vs. diabetic mice group.

### Human ASCs in mIslet+hASC scaffolds improve mouse islet function *in vivo*

The levels of circulating mouse specific C-peptide measured on day 60 post implantation showed a significant increase in mice implanted to the IP site with mIslet+hASC compared to the mIslet-alone group (Fig. 10a). The C-peptide levels were similar to the mice implanted with free-floating mouse islets under kidney capsule and control non-diabetic mice (Fig. 10a). Similarly, we found improvement in the ratio of mouse C-peptide to fasting blood glucose in mice implanted with mIslet+hASC scaffolds compared to mIslet-alone on day 60 post implantation (Fig. 10b). A control analysis of the mouse pancreas for presence of native insulin showed no mouse insulin in the mouse pancreas (Method supplement and Figure supplement 3). This confirms that the detected mouse C-peptide was secreted by the islets within the scaffolds. Explantation of the mIslet+hASC scaffolds and immunofluorescent staining of scaffolds for insulin and CD105, which is one of the ASC surface markers [48], confirms the presence of both islets and ASCs in the scaffolds post implantation (Fig. 10c).

**Fig 10.**
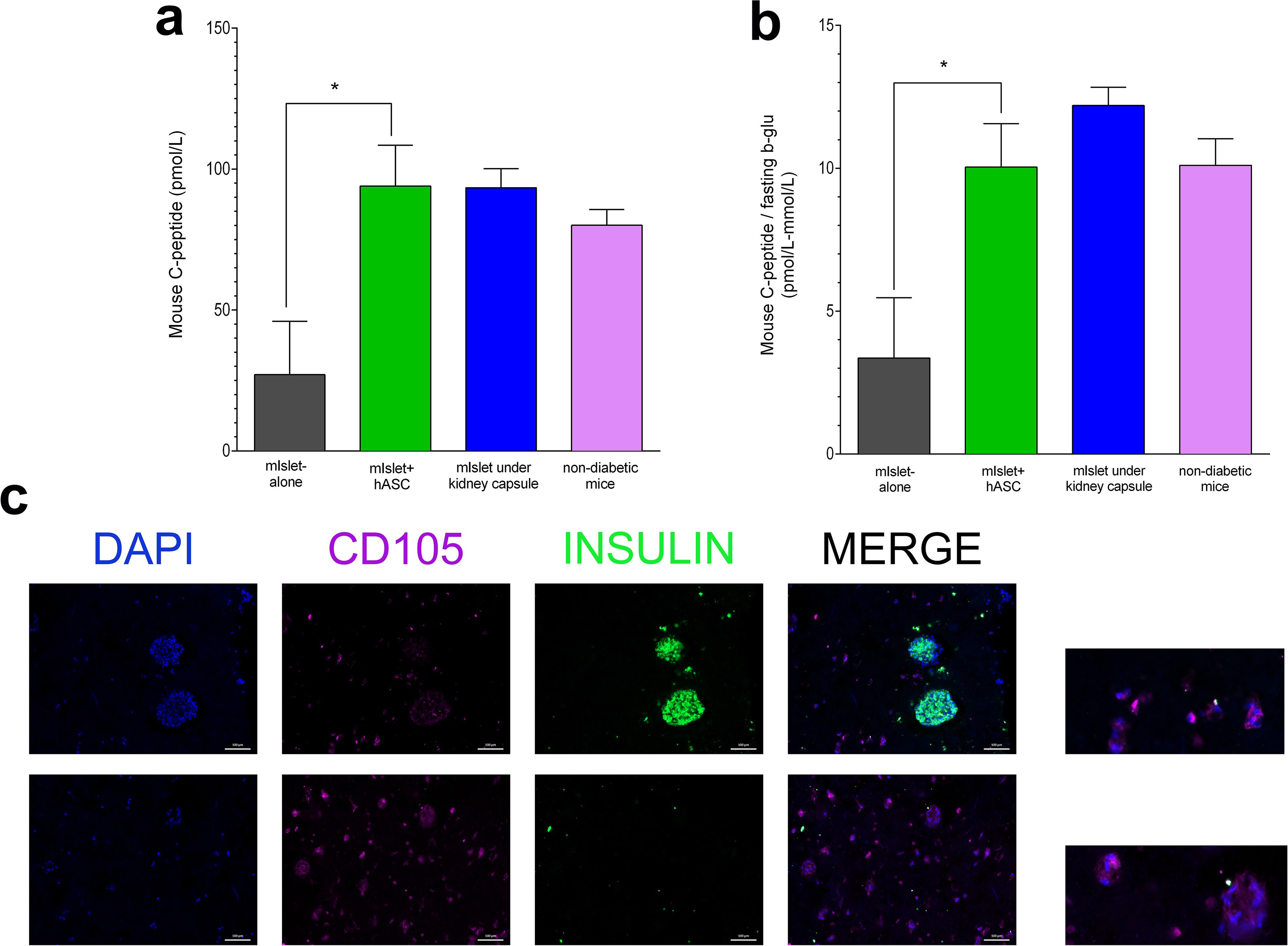
with 1 supplement (ESR Fig 3): 3D bioprinted mIslet+hASC scaffolds improve mouse islet function *in vivo*. Circulating mouse C-peptide (a) and calculated ratio of mouse C-peptide to fasting blood glucose (b) at the termination of studies for diabetic mice implanted with either scaffolds or mouse islet grafts under kidney capsule. Immunofluorescent double-staining of explanted scaffolds for insulin and CD105 (c) 30 days post implantation. Magnification 10x, scale bar 200 µm. In all analysis, data is presented as mean±SD and analysed by one-way ANOVA with Bonferroni corrections and unpaired Mann Whitney U-test. N=3 independent biological replicates and for each replicate 6 mice per group were used. * p<0.05 vs. mIslet-alone group.

## Discussion

Our results reveal the diffusion properties of alginate/NFC bioink to a wide-range of small molecules between 3-70kDa and we showed with micro-CT that we had an even distribution of islets in our 3D bioprinted scaffolds. Both islets and ASCs stayed viable throughout the studies in a common culture medium and ASCs in the double-layered islet scaffolds improved significantly both viability and function of the pancreatic islets. We report that ASCs decreased secretion of selected pro-inflammatory cytokines in both mouse and human islet scaffolds, which resulted in reduced cell death and improvement of islet function in the scaffolds. We also show the functional minimal mass of hIslet-alone scaffolds at the IP site of diabetic mice. Interestingly, our mIslet+hASC scaffolds were able to normalize blood glucose of diabetic mice 7 days post implantation at the IP site followed by increase in the levels of secreted C-peptide by the implanted scaffolds. Both islets and ASCs stayed viable throughout the *in vivo* studies as confirmed by immunofluorescent staining of explanted scaffolds for islet and ASC markers.

One of the first attempts to 3D bioprint human islets with collagen I gel supplemented with extracellular matrix (ECM) components, fibronectin and collagen IV showed preserved expression of pancreatic genes and insulin release profile similar to the freshly isolated human islets [49]. Although this study reported the printability of pancreatic islets, the islets stayed viable only up to 10 days post printing *in vitro* with no *in vivo* function. [49] Since then it has been well-known that choosing a right biomaterial for 3D bioprinting is one of the key factors that plays a vital role in islet function in a scaffold. Alginate with fast cross-linking ability and high water content creates quickly a robust structure that facilitates exchange of nutrients, oxygen and waste through the 3D bioprinted scaffolds [50]. However, alginate has poor printability and combining it with other hydrogels with higher fidelity such as gelatine has been reported to improve printability. However, scaffolds with either islets or INS1E beta cells showed viability and function only for 7 days post printing *in vitro* but no *in vivo* function [20, 29]. Combining alginate with NFC, which has the outstanding properties in shear thinning and printability, has been shown to have desirable properties for 3D bioprinted scaffolds of living cells [25, 51]. Liver cells and induced pluripotent stem cells (iPSCs) have been printed and maintained in alginate/NFC scaffolds [50, 52] and this combination was reported to be suitable for keeping the pluripotency of iPSCs that was later used to generate cartilage tissue [53]. In the recent studies, combination of tunicate-derived NFC with alginate was used to 3D bioprint human ear chondrocytes and autologous micro-fat for pre-clinical applications [13, 54]. Our data shows capabilities of this bioink mixture to diffuse wide-range of small molecules from 3 to70 kDa, which confirms its ability to exchange nutrients, waste and oxygen between the cells and outer environment.

Micro-CT imaging has been used previously to study the shape and the pore size of the 3D bioprinted scaffolds but not for studying the number of cell clusters in a scaffold [49]. To visualize the islets inside the bioprinted scaffolds, we used phosphotungstic acid (PTA), which enhances the contrast for the images taken by micro-CT [55]. Phase-contrast imaging using micro-CT with PTA staining has been shown to give high-quality visualization of chondrocytes in articular cartilage [55, 56]. Using this technique, we were able to monitor the number of islets per scaffold, which is a crucial step in generating functional 3D bioprinted islet scaffolds with good reproducibility.

Although, printability of pancreatic islets by using biocompatible and printable bioinks has been reported previously, the long-term function of these islet scaffolds has been hindered post printing. Creating a supportive micro-environment for the islets can have a direct impact on improving islet function and health. Combining bioink with ECM molecules and anti-inflammatory cytokines has been reported as a strategy to support islet function and viability. However, challenges such as the short half-life of the ECM molecules, growth factors and cytokines are considerable factors that affect the long-term outcome of these strategies [29, 49, 57, 58]. ASCs express various growth factors, cytokines and intracellular adhesion molecules [32]. The paracrine effects of ASCs for prolonging islet survival and function as well as reducing inflammatory reactions have been reported with the means of co-culture and co-transplantation of islets and ASCs [32, 37]. Direct co-culture of islets and ASCs induces adhesion of ASCs on the islet surface. This can change the morphology of an islet and affect the potency of insulin secretion. However, indirect culture of ASCs on a Petri dish and islets in transwells in close proximity has been reported to be beneficial through secretion of trophic factors from ASCs to improve islet function [59].

Previously, we have established harvesting and production of human ASCs from fat tissue according to good manufacturing practice (GMP) guidelines for clinical cell transplantation applications [60]. Here in this study, our data reveals that bioprinting of islets and ASCs in double-layered scaffolds improve islet viability and insulin secretion and support the previous findings on the beneficial effect of ASCs on islets in indirect culture conditions. In addition, reduction in secretion of the three measured pro-inflammatory cytokines IP-10, GRO-α and MCP-1, which all have been reported to increase in clinical islet transplantation in T1D patients, revealed the importance of ASCs as supporting cells to improve the islet health and consequently to elevate the outcome of islet transplantation [61–63]. We used an extrusion-based bioprinter that extrudes the bioink with pressurized air through the bioprinter nozzles. The printing pressure for both layers was between 5-12 kPa, which is tolerable for most cells types and we selected the size of the nozzles suitable for the size of islets and ASCs [25, 52]. However, the observed reduction in function and viability of islet-alone scaffolds could be due to the printing procedure and changes in islet micro-environment. This data on islet-alone scaffolds emphasizes the positive impact of adding ASCs to create supportive micro-environment for islets.

400 human islets is the minimum number of islets per graft under the kidney capsule to study the potency of pancreatic islets in a diabetic mouse model [64]. We show equal functionality of the minimal mass of human islets in 3D bioprinted scaffolds implanted to the IP site of diabetic mice compared to the minimal mass of islet grafts implanted under kidney capsule. We found no significant different in the levels of human C-peptide and the ratio of human C-peptide to fasting glucose in mice implanted with either scaffolds or free-floating islets under the kidney capsule. C-peptide to fasting blood glucose ratio has been known as the sign of preserved islet function post implantation [65]. The observed significant increase in the levels of human C-peptide in mice implanted with full mass of islet grafts (800 islets/graft) and the ability these islet grafts to reverse diabetes can be due to the higher number of islets per graft. This level of measured human C-peptide in our diabetic mice implanted with full mass of islet graft under kidney capsule has been shown previously to correlate with functional human islet graft in clinical settings [47]. Human islets are scarce material and being able to improve the clinical outcome of islet transplantation with minimal islet mass in scaffolds would benefit many T1D patients. Using ASCs as supporting cells together with islets could be a revolutionary strategy for having functional islet scaffolds for clinical applications.

In order to test this strategy and due to the lack of readily available human islets for this study, we fabricated 3D bioprinted scaffolds with 100 mouse islets and 1.2×10^6^ human ASCs. The implantation of mIslet+hASC scaffolds to the IP site of diabetic mice could normalize blood glucose 7 days after implantation and this is almost similar to implanting full mass of mouse islets per graft (250 islets) under kidney capsule of diabetic mice. Similar levels of secreted mouse C-peptide and the ratio of mouse C-peptide to fasting blood glucose were observed in mice that received mIslet+hASC scaffolds to the IP site vs. full mass mouse islet grafts under kidney capsule. The implantation of a scaffold to the IP site of the diabetic mouse model is beneficial due to the easy accessibility and the larger space for different shaped scaffolds as these strategies are developing towards clinical applications [66]. However, delayed delivery of oxygen and nutrient supplies due to the passive diffusion properties compared to the kidney capsule site could hinder the outcome of transplanted scaffolds to the IP site [67]. Our data shows the equal levels of functionality for mIslet+hASC scaffolds at the IP site vs. full mass of mouse islet grafts under kidney capsule of diabetic mice. Explanted mIslet+hASC scaffolds were positively-labelled for islet and ASC markers, which confirm the presence of ASCs in the scaffolds together with islets throughout the *in vivo* studies. Five-times Intravenous administration of 5×10^6^ ASCs to diabetic mouse models has been shown less efficient since ASCs have been detected in lungs, spleen and peritubular regions but not in the pancreas post infusion [68]. However, our data shows the presence of ASCs in scaffolds 60 days post implantation and this can support the beneficial effects of ASCs on preserving the islet function. Our data suggests the possibility of having a functional islet scaffold with only 100 mouse islets per scaffold when they are combined with ASCs.

One of the limitations with performing transplantation studies with mouse islets to diabetic mouse models is the chance of native mouse beta cell regeneration within the pancreas post chemical beta cell ablation [69]. Previously, islet regeneration in mouse pancreas has been reported more than 90 days post islet ablation [69]. We found no detectable insulin from the mouse pancreas 60 days post implantation. This confirms the contribution of the 3D bioprinted scaffolds in normalizing the mouse blood glucose. Another alternative strategy would be to remove implanted scaffolds and observe the reoccurrence of the diabetes in a mouse model [70]. However, this strategy would require another operation for the animals with a follow-up afterwards, which could affect negatively the quality of the studies *in vivo* and animal welfare.

This study presents fabrication of a double-layered device for delivery of pancreatic islets and ASCs by using alginate/NFC bioink generated by means of 3D bioprinting technology. Our scaffold demonstrates the paracrine effects of ASCs to prolong islet function and viability. Our data on a long-term functional islet+ASC scaffold implanted at the extrahepatic site of diabetic mouse model opens a window of opportunities for creating a fully functional transplantable islet scaffold. Next generation diabetes treatment with the means of beta cell replacement therapy relies on generation of functional islet scaffolds that can be implanted to the extrahepatic sites as well as recreation of physiological micro-environment for pancreatic islets or stem cell-derived beta cells. Our scaffold is an important step toward an improved strategy for fabricating translatable, safe and fully functional 3D bioprinted islet structures for clinical applications.

## Materials and Methods

### Ethics

All experiments and methods using human islets were approved by and performed in accordance with the guidelines and regulations made by the regional committee for medical and health research ethics central in Norway (2011/782). The animal experiments were approved by the Norwegian National Animal Research Authority (permit number FOTS ID 10680) and performed according to the Guide for the Care and Use of Laboratory Animals published by the US National Institutes of Health (NIH Publication, 8th Edition, 2011), and to the Norwegian Animal Welfare Act. All animals were handled by an experienced animal technician at all times and blinded as the technician was not aware of the difference between groups. Regarding the use of ASCs, all donors signed informed consent, and the use of ASCs was approved by the Regional Committee for Medical and Health Research Ethics (2014/838).

### Cell isolation and maintenance

#### Human pancreatic islets

Human islets from non-diabetic donors were obtained from the JDRF award 31-2008-416 (ECIT Islet for Basic Research program) and isolated as previously described [71] from male/female 4/3 non-diabetic brain-dead donors with mean age 53 years (46–59 years) and mean BMI 20 (23–29 kg/m^2^) after appropriate informed consent from relatives for multi-organ donation and for use in research. Islet purity of >80% was used in this study, judged by digital imaging analysis [72] or dithizone staining. Equally sized islets were manually hand-picked and distributed blindly among experimental groups. Islets were cultured at 37°C (5% CO2) up to 48 hrs on petri dishes (Sterilin, Newport, UK) with CMRL 1066 medium supplemented with 10% human AB serum (Milan ANALYTICA, Rheinfelden, Switzerland), 1% penicillin/streptomycin, 10 mmol/L HEPES (Life Technologies, Carlsbad, CA, USA) prior to start of the experiments.

#### Mouse pancreatic islets

Mouse islets were isolated from 8–18 weeks old male Balb /c Rag 1^-/-^ mice (Taconic, Denmark) as previously described [73]. See Method supplement for more details.

#### Human ASCs

Lipoaspirates were obtained from the flank or thigh of six females aged 60.4 ± 4.2 years undergoing elective plastic surgical operations at the Department for Plastic Surgery, Radiumhospitalet, Oslo University Hospital. The stromal vascular fraction (SVF) was extracted using the Celution® system (Cytori Therapeutics Inc., San Diego, CA, USA) according to the manufacturer’s instructions. See Method supplement for more details [60].

### Preparation of the bioink and 3D bioprinting of the scaffold

The double-layered scaffold with the total size of 10 × 10 × 1 mm was designed with TinkerCAD and the stl file was converted with Slic3r slicing program into G-code and manually adjusted to its final format. Commercially available CELLINK^®^ Bioink (CELLINK AB), which is a plant-derived NFC and alginate was used for bioprinting the cell scaffolds [26].

For the bioprinting, ASCs were mixed with the bioink between two locked syringes using a female/female luer lock adapter (CELLINK AB). After mixing, the bioink-ASC mixture was loaded to the printer cartridge and connected with a blue standard conical 22G (410 μm) nozzle for bioprinting the bottom solid layer of the scaffold. 1.2 × 10^6^ ASCs were used per scaffold. The top islet layer in shape of orthogonal grid of filaments was bioprinted with a pink standard conical 20G (580 μm) nozzle to avoid excessive shear stress on the islets. Islets were hand-picked and mixed with the bioink on a petri dish gently with a spatula. 100 islets/scaffold were used for all *in vitro* analysis as well as *in vivo* studies with mouse islets and 400 islets/scaffold were used for *in vivo* studies with human islets. The scaffolds were bioprinted with pneumatic extrusion bioprinter INKREDIBLE+ (CELLINK AB) with the printing pressure of approximately 5-12 kPa for both ASC and islet layers. Printed scaffolds were crosslinked with 50 µM CaCl_2_ CELLINK Crosslinking Agent (CELLINK AB) for 5 min. mIslet-alone, mIslet+hASC, hIslet-alone or hIslet+hASC were all cultured in MEM-α culture medium complemented with 10% human platelet lysate (HPL), 1% penicillin/streptomycin, 10 mmol/l HEPES (Life Technologies, Carlsbad, CA, USA) for further analysis.

### Micro-CT

Islets were stained with 0.3% PTA aqueous solution overnight at room temperature. The islets were then rinsed 2 times in 1x PBS followed by mixing the islets with the bioink and bioprinting of the islet scaffolds using the same process described in section for preparation of the bioink and 3D bioprinting. See Method supplement for more details.

### Fluorescence Recovery After Photobleaching (FRAP)

Circular NFC/alginate discs (Ø 20 mm) were designed, printed and crosslinked similarly as described above. Printed discs were incubated in 0.1 mg/mL FITC-labelled dextran molecules (size either 3-5 kDa, 10 kDa, 20 kDa or 70 kDa (Sigma-Aldrich)) dissolved in PBS with 20mM CaCl2 (Sigma) overnight. FRAP was performed on 5 different spots per hydrogel disc with a Leica TCS SP8 STED confocal microscope with an Ø60 μm bleaching area and frame rate of 0.223 s for 120 s. Fluorescence recovery curves were obtained through image analysis with the open source software FIJI (https://fiji.sc/). The time required for a bleach spot to recover half way of the recovery curve (τ1/2) was determined through FRAPbot (http://frapbot.kohze.com/). Apparent diffusion constants (D) were determined based on the theory of Soumpasis et al, [74, 75] utilizing formula 1, with r the radius of the bleach laser.

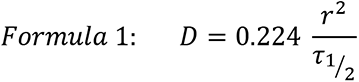

### Glucose-stimulated insulin secretion assay

Human or mouse islet scaffolds w/wo ASCs were incubated in 1.67 mmol/l glucose for 2 hrs at 37°C followed by 2 hrs incubation with 20 mmol/l glucose and 2 hrs incubation with 1.67 mmol/l glucose. Insulin secretion was analysed in the supernatant fraction using human-specific or mouse-specific insulin ELISA (Mercodia, Uppsala, Sweden). See Method supplement for details.

### Viability analysis of the cells in 3D printed scaffolds

Viability assessment was performed on mouse or human islet scaffolds w/wo ASCs using FDA/PI imaging to evaluate live vs. dead cells in scaffolds on days 1, 8, 14 post printing. The level of Adenylate Kinase (ADK) enzyme indicating cell toxicity was measured in the culture medium harvested on days 1, 8 and 14 post printing according to the manufacturer’s protocol (Lonza, Basal, Switzerland). See Method supplement for details

### Biochemical measurements

The level of mouse and human MCP-1, IP-10 and GRO-α cytokines were measured in culture medium of islet scaffolds w/wo ASCs on days 1, 8 and 14 post print using multiplex immunoassay according to the manufacturer’s protocol (Thermo Fisher, MA, USA).

### *In vivo* implantation of scaffolds

The animals were housed no more than 5 mice per cage and were maintained in a 12h light/dark cycle in an approved unit with free access to food and water except during fasting. Prior to implantation experiments, diabetes was introduced to 8-10 week-old male Balb/c Rag 1^−/−^ immunodeficient mice (C.129S7(B6)-Rag1tm1Mom/J, stock 003145, The Jackson Laboratory, Sacramento, California, USA) by administration of one dose of Alloxan monohydrate 75 mg/kg intravenously (Sigma Aldrich, Oslo, Norway) 3 days prior to the islet implantation. Mice with non-fasting blood glucose ≥20 mmol/l for 2 consecutive days were selected as diabetic recipients. For *in vivo* studies with mouse islets, scaffolds containing 100 Balb/c Rag 1^−/−^ mouse islets w/wo human ASCs were implanted at the IP site of the diabetic mice. Along with the mice received mouse scaffolds w/wo ASCs, 250 free-floating Balb/c Rag 1^−/−^ mouse islets of the same islet batch were hand-picked and implanted under the left kidney capsule as described previously [73]. N= 3 independent biological replicates were performed. 6 mice were used in each of the mIslet+hASC and mIslet-alone groups, 2-3 mice were used in each of the diabetic, non-diabetic and mIslets under kidney capsule groups.

For *in vivo* studies with human islets, scaffolds containing 400 human islets, minimal islet mass in scaffold (hIslet-alone minimal mass) were implanted at the IP site of the diabetic mice. In parallel, 800 free-floating human islets (full mass under the kidney capsule) and 400 free-floating human islets (minimal mass under kidney capsule) were also implanted under the left kidney capsule of diabetic mice. N= 3 independent biological replicates were performed. 6 mice were used in each of the groups. 2-3 mice were used in each of the diabetic and non-diabetic groups.

Random non-fasting blood glucose was monitored and measured every 2 to 3 days until the termination of the studies. *In vivo* studies with mouse islets were terminated on day 60 post scaffold/islet implantation. *In vivo* studies with human islets were terminated on day 30 post implantation of scaffolds/islets. Fasting blood glucose was measured throughout the study. At the termination of studies, mice under anaesthesia were sacrificed by heart puncture for blood samples. Plasma samples were prepared for further analysis of mouse-specific and human-specific C-peptide. Mouse pancreases from mouse studies were harvested and stored in liquid nitrogen for further analysis of native mouse-specific insulin (Mercodia, Uppsala, Sweden). See Method supplement for more details.

### Immunofluorescent staining and imaging of the scaffolds

At the termination of the *in vivo* studies, scaffolds were explanted and fixed in 4% paraformaldehyde and embedded on O.C.T. compound for further immunofluorescent staining analysis for insulin, C-peptide, glucagon and CD105. Images were taken with Axio Observer Inverted Microscope (Carl Ziess AS) operated by ZEN lite software. 5 representative images were taken per scaffold. See Method supplement for more details.

### Statistical analysis

Data are presented as means ± SD. Differences among the 3 groups were evaluated by one-way ANOVA with Bonferroni corrections. A Mann–Whitney U test was performed for difference analysis between 2 groups. Significance was set at p < 0.05. Data were analysed using GraphPad Prism software, version 6.0 (La Jolla, CA, USA)

## Acknowledgements

The authors are grateful to all members at the human islet isolation facility at the Oslo University Hospital, Oslo, Norway. We especially thank Merete Høyem, Marina Katavic and Ragnhild Fjukstad for technical expertise and helpful discussions. This work was supported by the Research Council of Norway through its Centres of Excellence funding scheme, project number 262613, The Norwegian Diabetes Association, Oslo Diabetes Research Center, Southern and Eastern Norway Regional Health Authority, project number 2019027 as well as Research Council of Norway, project number IPN 317790.

## Author contributions

S.A. and H.S. conceived, designed and performed the experiments, and wrote the manuscript. S.A., E.M.N., L.S.O., C.H., R.V., L.P.N. performed experiments, and analysed data. P.G. and A.A. provided expertise, feedback, and designed experiments. H.S., P.G. and S.K. secured funding. S.A. and H.S. performed human islet isolation. All authors contributed to manuscript revision, read, and approved the submitted version. S.A and H.S are the guarantors of this work, and as such have full access to all data in the study and take responsibility for the integrity of the data and the accuracy of the data analysis.

## Data availability

All data that support the finding of this study are openly available in the Dryad repository (https://datadryad.org/stash/share/SgvW5qQJik0-1Kx9vHe64LrIWi1s5H93cCP7_3wjoMw). The corresponding authors confirm that the data supporting the findings of this study are also available within the article and its supplementary materials.

## Corresponding authors

Correspondence and request of materials should be addressed to Shadab Abadpour and Hanne Scholz.

## Appendix 1

### Method supplement

#### Mouse pancreatic islets

Mouse islet isolation procedure were performed by using 3.0 ml of Hank’s balanced salt solution containing 8mg/ml Collagenase P from Clostridium histolyticum (Roche, Mannheim, Germany) injecting into the pancreatic duct. The distended pancreas was subsequently removed and incubated at 37°C for 17 min followed by gradient purification of the endocrine tissue. The islets were placed in 90-mm-Petri dishes (Sterilin, Heger AS, Norway), and cultured overnight in RPMI 1640 media without L-glutamine (HyClone, Utah, USA) supplemented with 10% heat-inactivated fetal bovine serum, 1% penicillin/streptomycin, 10 mM 5ml Hepes and 1% L-glutamine (Gibco, Paisley, UK) at 37°C (5% CO2) until use in experiments.

#### Isolation and Culture of ASCs

The adipose tissue obtained through liposuction was washed with Ringer’s solution and enzymatically digested by a proprietary enzyme solution (Celase®) during constant agitation, followed by washing and concentrating of the cells using centrifugation. Directly after processing, the SVF was washed once with 5% human serum albumin (Octapharma, Jessheim, Norway) and centrifuged at 600 × g for 10 min at 21°C. Mononuclear cells were counted using a hemocytometer (Kova, Garden Grove, CA, USA) and seeded at 3,000 cells/cm2 in a T75 flask (Nunc; Thermo Fisher Scientific, Waltham, MA, USA) with supplemented minimum essential medium MEM-α containing 10% fetal bovine serum (FBS) (both from Gibco, Thermo Fisher Scientific) and 50 μg/ml gentamicin (Braun, Esbjerg, Denmark). Cells were cultured at standard condition of 37°C in a humidified atmosphere with 21% O2 and 5% CO2. The primary cells were allowed to attach for 2 days before being washed 3 times with 37°C phosphate-buffered saline (PBS; Lonza, Basel Switzerland) to remove nonadherent cells. The culture medium was changed every 2–3 days until the cells reached 70%–80% confluency, followed by harvest of the cels using TrypLE Express (Gibco). Harvested vcells were subsequently seeded at a density of 3,000 cells/cm2 for the next passage. ASCs of passages 1-2 were cryopreserved in medium containing 10% dimethyl sulfoxide (DMSO; Cryo-Sure; Wak-chemie, Steinbach, Germany) and 20% human serum albumin and initially cooled to −80°C at a rate close to −1°C/min using a freezing container (Mr. Frosty; Thermo Fisher Scientific). This is followed by transferring the cells to a −196°C liquid nitrogen vapor tank for storage. Cells were thawed in room temperature (RT), and DMSO was rapidly diluted in supplemented ASC medium before centrifugation and use in experiments.

#### Micro-Computed Tomography (Micro-CT)

In order to perform micro-CT analysis on bioprinted islet scaffolds, the scaffolds were placed into an eppendorf tube for scanning at a micro-CT system Bruker 1172 (Kontich, Belgium). Samples were scanned at 55 kV and 170 µA, with an exposure time of 250 ms per projection and frame averaging of 4, making a total of 1000 ms per projection. A total of 539 projections were acquired around 180°+ at a field of view of 1100 x 1332 pixels (W x H). The datasets were reconstructed using NRecon (Bruker microCT version 1.7.5.9), and further segmentation and analysis performed using Dragonfly software (version 2020.2, Object Research Systems (ORS) Inc, Montreal, Canada).

For the analysis, the samples were segmented using Otsu [1] to separate the islets from the constructs and background. The islets were individually labelled and further quantifications were performed using Dragonfly. The analysed parameters were the number of islets, the volume of each individual islet, the local thickness of the islets (calculated as the diameter of a hypothetical sphere that fits within each surface point of the islet) [2], sphericity [3] and length and width. The length and width were calculated based on the Feret Diameters, which are the measurements of an object size along a specified direction. This can be defined as the distance between the two parallel planes restricting the object perpendicular to that direction [4]. The length is calculated by taking the perpendicular distance between two parallel tangent planes touching the particle surface, after sampling 400 different orientations to find the largest distance between these two planes. The width is calculated by taking the two planes perpendicular to those two first ones and enclosing the object outline. 3 dimensional (3D) images were rendered using Dragonfly and colour-coded based on the local thickness of the islets.

#### Glucose-Stimulated Insulin Secretion assay (GSIS)

Day 1, 8 and 14 post printing, human or mouse islet scaffolds with/without (w/wo) ASCs were incubated for 2 hrs at 37°C in Krebs-Ringer bicarbonate buffer (11.5 mmol/l NaCl2, 0.5 mmol/l KCl, 2.4 mmol/l NaHCO3, 2.2 mmol/l CaCl2, 1 mmol/l MgCl2, 20 mmol/l HEPES, and 2 mg/l albumin: all Sigma-Aldrich, Oslo Norway) supplemented with 1.67 mmol/l glucose (low glucose). This is followed by 2 hrs incubation with Krebs buffer containing 20 mmol/l glucose (high glucose) and 2 hrs incubation with Krebs buffer containing low glucose. To analyze the mouse or human islet quality post isolation, twenty free-floating equally-sized mouse or human islets were handpicked and transferred into a transwell plate (Corning, NY, USA). Islet cells were incubated with krebs buffer supplemented with low glucose levels for 45 min at 37°C followed by incubating the cells with high glucose and then low glucose for 45 min in each solutions. Insulin secretion was analysed in the supernatant fraction using human- or mouse-specific insulin enzyme immunoassay (EIA) (Mercodia, Uppsala, Sweden). Stimulation index (SI) was calculated as a ratio of insulin secreted at high glucose (20 mmol/l) versus low glucose (1.67 mmol/l).

#### Viability analysis of cells in 3D printed scaffolds

Assessment of cell viability on human or mouse islet scaffolds w/wo ASCs were performed on days 1, 8 and 14 post print using fluorescein diacetate (FDA) 20µg/mL (Sigma-Aldrich, Oslo, Norway) for detection of live cells and propidium iodide (PI) 100µg/mL (Thermo Fisher Scientific) for evaluating the degree of dead cells. 20mM CaCl_2_ were added to all solutions to avoid degradation of scaffolds. This is followed by imaging of the scaffolds using Axio Observer Inverted Microscope (Carl Ziess AS) operated by ZEN lite software. 5 images / scaffold were taken representing each scaffold on the indicated time-points.

The level of cytoxicity was analyzed by measuring adenylate kinase (ADK) enzyme in the culture medium of mouse or human islet scaffolds w/wo ASC harvested at days 1, 8 and 14 post printing. The measurement was performed by using the toxilight non-destructive cytotoxicity bioassay kit according to the manufacturer’s protocol (Lonza, Basal, Switzerland).

#### Analysis of native mouse insulin in pancrease of mice transplanted with scaffolds containing mouse islets w/wo ASCs

At termination of *in vivo* mouse studies on day 60 post implantation, pancreas were harvested and stored in liquid nitrogel. Frozen pancreas samples were crushed by using mortar and pestle. Tissue lysis buffer (RIPA buffer containing halt protease-phosphatase inhibitors (Thermo scientific, Oslo, Norway) was added to the pancreas tissue samples proceeding to mechanical disruption using sonication. Samples were centrifuged and purified using QIAshredder purification column (QIAGEN, Hilden, Germany). Total protein concentrations were determined using Pierce BCA protein assay (Life Technologies AS, Oslo, Norway). To measure the total insulin in the lysate, acetic-ethanol was added to the lysates and incubated at 4°C for 24 h before insulin content was measured by human insulin ELISA kit (Mercodia AB) and presented as insulin content over total protein concentration of samples.

#### Immunofluorescent Staining and imaging of scaffolds

Fixated alginate/NFC scaffolds were incubated in 15% sucrose solutions overnight in 4 °C, followed by the overnight incubation in 30% sucrose and 60% sucrose/OCT solutions respectively. 20mM CaCl_2_ were added to all solutions. The scaffolds were placed in OCT and snap freeze with liquid nitrogen. Scaffold slides were stained with guinea pig anti-human insulin polyclonal antibody 1:500 (DAKO, Cat# A0564, RRID:AB_10013624, Oslo, Norway), rabbit anti-human glucagon polyclonal antibody 1:100 (DAKO, Cat# A0565, RRID:AB_10689823, Oslo, Norway), mouse anti-human CD105 1:100 (Thermo Fisher Scientific Cat# MA5-11854, RRID:AB_10985671, Oslo, Norway) and guinea pig anti-human C-peptide 1:100 (Abcam, Cat# ab30477, RRID:AB_726924, Cambridge, UK). Donkey anti-mouse Alexafluor 594 (Thermo Fisher Scientific Cat# A-21203, RRID:AB_141633), goat anti-guinea pig Alexafluor 488 (Thermo Fisher Scientific Cat# A-11073, RRID:AB_2534117) and goat anti-rabbit Alexafluor 594 (Thermo Fisher Scientific Cat# A-11012, RRID:AB_2534079), all in dilution of 1:300 (Oslo, Norway) were used as secondary antibodies to detect expression of antibodies followed by nuclear staining using SlowFade Gold antifade reagent with DAPI (Life Technologies AS, Oslo, Norway). Images were taken with Axio Observer Inverted Microscope (Carl Ziess AS) operated by ZEN lite software. 5 representative images were taken per scaffold.

### Result supplement

#### FRAP reveled diffusion of small molecules with various sizes to alginate/NFC bioink

Photobleaching of a small area of our bioink by using FRAP analysis showed diffusion of molecules with sizes 3-5 kDa (Video supplement 1), 10 kDa (Video supplement 2) and 70 kDa (Video supplement 3) to the bioink and full recovery of the bleached area.

#### Imaging and calculation of human islets in 3D bioprinted scaffolds used for *in vitro* analysis of human islet function and viability post printing

Using micro-CT imaging technique, human islets within 3D bioprinted scaffolds were imaging. Micro-CT analysis showed an even distribution of human islets within 3D bioprinted scaffolds (Figure supplement 1, Video supplement 4).

#### Imaging and calculation of human islets in 3D bioprinted scaffolds used for *in vivo* analysis of human islet function post printing

Using micro-CT imaging technique, human islets within 3D bioprinted scaffolds were imaging. Micro-CT analysis showed an even distribution of human islets within 3D bioprinted scaffolds (Figure supplement 2, Video supplement 5).

#### No detection of native insulin in the pancreas of mice implanted with islet scaffolds w/wo ASCs as well as free-floating mouse islets under kidney capsule

The level of native mouse insulin was measured in the pancreas of mice implanted with scaffolds containing mIslet+hASC and mIslet-alone. In addition, this insulin level was also measured in mice transplanted with free-floating mouse islets under kidney capsule as well as non-diabetic healthy animals. Calculating the ratio of total mouse insulin to total protein in the pancreas showed no insulin detected in the pancreas of mice reiceved either islet scaffolds or islet graft (Figure supplement 3). However, native mouse insulin was detected in the pancreas of the non-diabetic healthy mice.

## Figure supplement legend

**Figure supplement 1:**
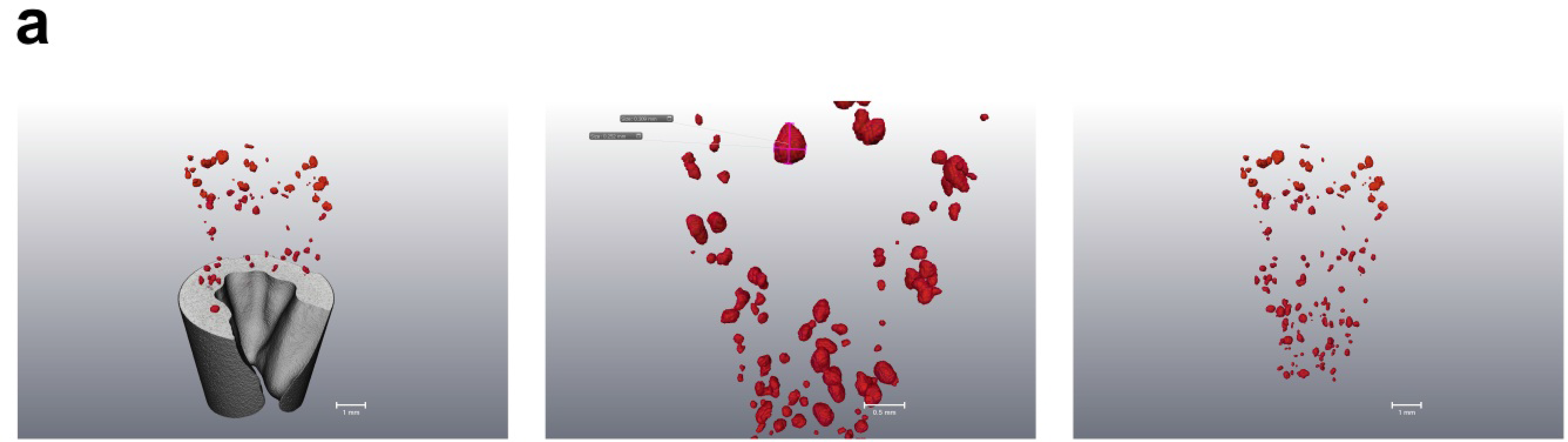
even distribution of 100 human islets per scaffold. Micro-CT analysis showing human islets in 3D bioprinted scaffolds.

**Figure supplement 2:**
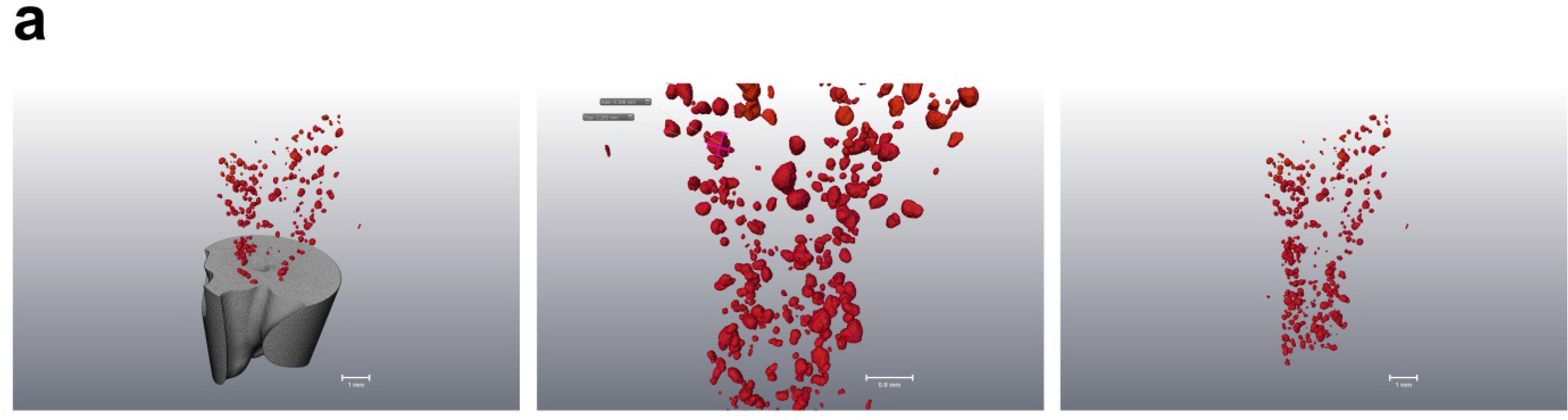
even distribution of 400 human islets per scaffold. Micro-CT analysis showing human islets in 3D bioprinted scaffolds.

**Figure supplement 3:**
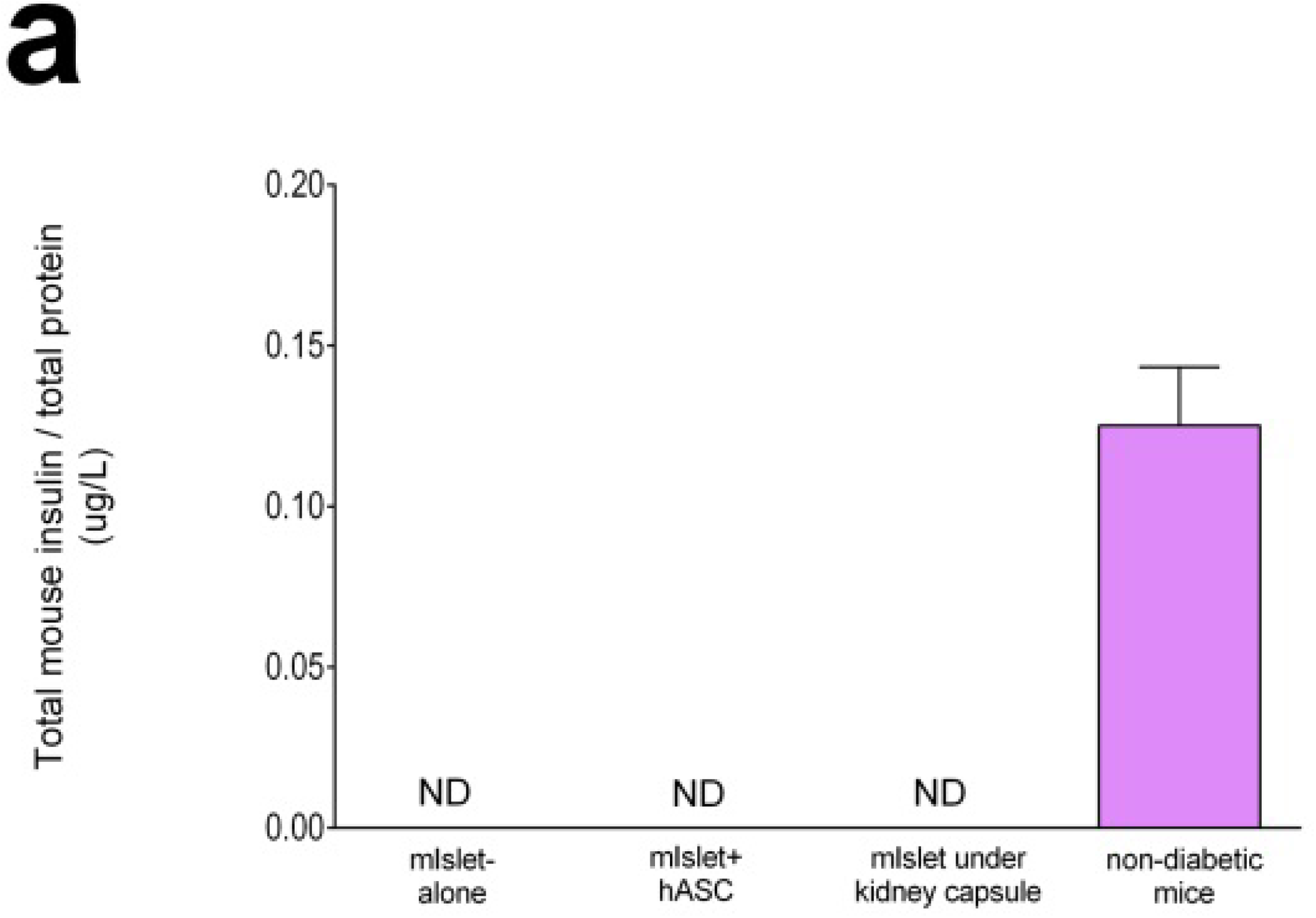
Measurement of native insulin in pancreas of mice received either scaffolds or islet grafts. The ratio of total insulin to total protein in pancreas was calculated in mice received mIslet-alone, mIslet+hASC, mIslet under kidney capsule as well as non-diabetic mice.

## Table supplement

**Table supplement 1: Micro-CT analysis revealed total number of islets per 3D bioprinted scaffold used for *in vitro* analysis of human islet function and viability post printing**

**Table supplement 2: Micro-CT analysis revealed total number of islets per 3D bioprinted scaffold used for *in vivo* analysis of human islet function post printing**

## Abbreviations

3D: 3 Dimensional
NFC: Nanofibrillated Cellulose
ASC: Adipose-derived Stromal Cell
Micro-CT: Micro-Computated Tomography
IP: Intraperitoneal
FRAP: Fluorescence Recovery After Photobleaching
IP-10: Interferon gamma-induced protein-10
MCP-1: Monocyte Chemoattractant Protein-1
GRO-α: Growth-regulated oncogene-α

